# Transcriptomic and proteomic responses to gas vesicle collapse in native and engineered bacterial systems

**DOI:** 10.64898/2026.05.31.729101

**Authors:** Manuel Iburg, Aaron O. Bailey, Art He, William K. Russell, George J. Lu

## Abstract

Gas vesicles (GVs) are air-filled protein nanostructures produced by microbes to regulate buoyancy and have emerged as powerful tools in biomedical imaging, particularly as acoustic reporter genes. Their mechanically robust shells enclose a stable air compartment, which collapses irreversibly when subjected to sufficient hydrostatic or acoustic pressure, leaving behind large protein sheets. This collapse phenomenon underlies key applications such as differential imaging and controlled cavitation and is also believed to occur naturally during buoyancy regulation, yet its physiological consequences remain poorly understood. Here, we used transcriptomic and proteomic approaches to investigate cellular responses to GV collapse. In the native GV-producing cyanobacterium *Dolichospermum flos-aquae*, RNA sequencing revealed a distinct transcriptional response characterized by the upregulation of heat shock proteins, indicative of a stress reaction to intracellular protein aggregation. In contrast, bioluminescence reporter assays in *E. coli* heterologously expressing GVs showed no comparable activation of heat shock promoters. To test the hypothesis that collapsed GVs can be recognized by specific proteins inside cells, we conducted LC-MS/MS-based pull-down assays in both species but did not identify strong candidate binders. While no definitive recognition mechanism was uncovered, our omics-based study provides a rich dataset for cellular responses to GV collapse in both the native and heterologous systems. These findings suggest that cellular responses to collapsed GVs may be more complex than previously recognized, and that both improved assay sensitivity and additional focused experiments will be needed to elucidate how cells detect and manage large intracellular protein aggregates such as collapsed gas vesicles. Proteomics data are available via ProteomeXchange (PXD060779), and RNA sequencing data via NCBI GEO (GSE289028).

## INTRODUCTION

Gas vesicles (GVs) are genetically encoded protein nanostructures that encapsulate ambient air^1-3^. They are natively found in microbes that express them to gain buoyancy, which enables photosynthetic microbes such as *Dolichospermum flos-aquae* (also known as *Anabaena flos-aquae*) to move vertically in a body of water, competing for optimal access to sunlight^4, 5^. GVs have been the focus of recent advances in biomedical research. Most notably, GVs are known as the acoustic reporter genes (ARGs)^6^ that have catalyzed breakthroughs in gene expression imaging in tumor and tumor-homing microbes^7-11^, nanoparticle imaging in tissues and therapeutic cells^12-15^, and biosensing^16-18^, among other imaging modalities^19-21^. Beyond imaging, an expanding suite of GV-based techniques has broadened their utility into payload delivery^22^, cellular manipulation^23, 24^, gas delivery^25, 26^, neuromodulation^27^, cell tracking^15, 28^, and pressure sensing^29, 30^. A defining physical property of GVs is their susceptibility to collapse when exposed to hydrostatic or acoustic pressure above a genotype-specific threshold, resulting in irreversible rupture of the protein shell and release of the internal gas. This collapse is exploited in imaging to enhance contrast by selectively removing GV signal^6, 31, 32^, and in actuation, where collapse-induced cavitation enables controlled release of cellular contents^13, 33^.

Although collapsing GVs inside cells is critical for many biomedical applications, the physiological consequences of this collapse and the fate of the resulting broken GV protein shell remain unclear. Transmission electron microscopy images of purified GV particles have shown that collapsed GVs retain largely intact pieces of protein shells^32^. This is not a surprise because the major shell proteins of GVs are tightly associated through dense hydrogen bonding, particularly the amide backbone hydrogen bonding in the central β-sheet region of the protein. These large, β-sheet-rich protein shells of GVs often exceed 100 nm in diameter and contain on the order of 10,000 copies of the amphipathic GV shell protein, and it is intriguing to consider how cells might degrade such large and stable assemblies^34, 35^.

Here, we investigated whether GV collapse elicits a physiological response in host cells. In the native GV-producing cyanobacterium *D. flos-aquae*, transcriptomic analysis using RNAseq revealed a distinct upregulation of heat shock proteins following GV collapse, consistent with a stress response to intracellular protein aggregation. In contrast, we used bioluminescence reporter assays to probe several heat shock promoters in *E. coli* heterologously expressing GVs, but did not observe a comparable response above our assay’s detection threshold. In the second part of this study, we tested the hypothesis that cells may possess a protein-based mechanism to recognize collapsed GVs. Using LC-MS/MS-based pull-down assays in both species, we searched for proteins that bind selectively to collapsed GVs. Although we did not identify strong candidate effectors, several protein hits emerged that warrant further functional investigation and highlight the need for more sensitive tools and continued efforts to understand GV-related cellular signaling.

## RESULTS

### GV collapse in *Dolichospermum flos-aquae* induces upregulation of molecular chaperones

We selected *D. flos-aquae* for this study because it is a native GV-producing species, in which the majority of cells naturally express high levels of GVs (**Figure 1A**). To evaluate the impact of GV collapse on host cells, we investigated the transcriptional response in *D. flos-aquae*. Cultures were subjected to water bath sonication to induce GV collapse, with untreated cultures serving as controls. Phase-contrast microscopy confirmed successful GV collapse in the treated samples (**Figure 1B**). Following treatment, total RNA was extracted and subjected to RNA sequencing (RNAseq) to enable a differential whole-transcriptome analysis between cells with intact and collapsed GVs (**Figure 1C**). RNAseq analysis revealed a distinct set of transcripts that were significantly upregulated in treated cells (**Figure 1D & Supplemental Table 1**), with molecular chaperones being prominently represented. Among these, *clpB*, an Hsp100 family protein known for its role in protein disaggregation^36^, was notably induced. It is worth noting that the genome of *D. flos-aquae* contains two homologs of *clpB*, as previously reported in other cyanobacterial species^37^, with *clpBII* specifically upregulated in our dataset. Additional heat shock proteins, including *HtpG, DnaK, DnaJ-like*, and *HtpX-like*, were also significantly enriched. Together, these findings support the existence of a specific transcriptional response to GV collapse in *D. flos-aquae*, involving the induction of genes associated with protein folding and cellular stress mitigation.

**Fig 1.**
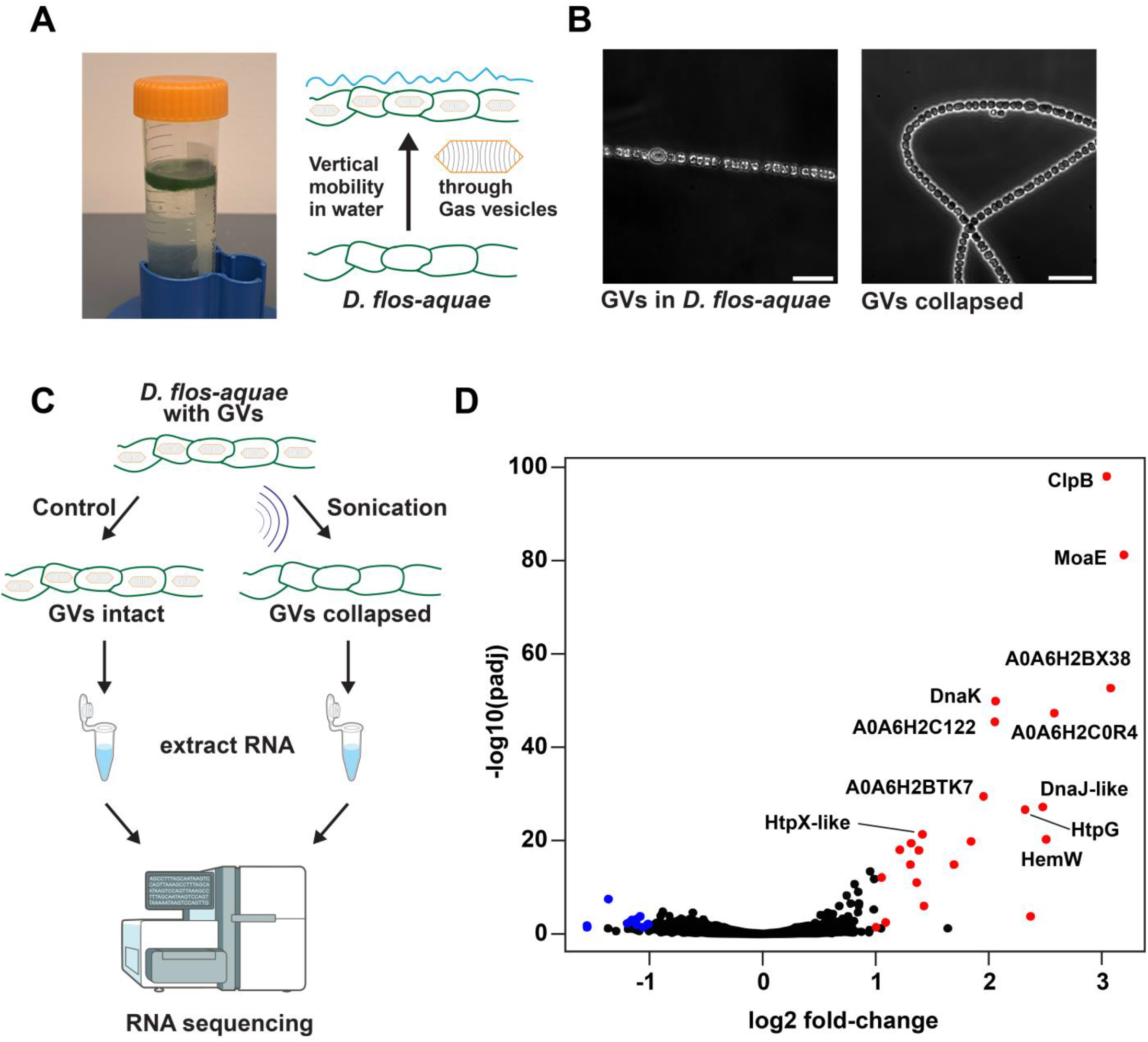
Distinct transcriptional response to gas vesicle (GV) collapse in *D. flos-aquae.* **(A)** Photograph of *D. flos-aquae* cultures floating in a 50 mL vessel due to intracellular GVs. Schematic illustrates how GVs confer buoyancy. **(B)** Phase contrast images of *D. flos-aquae* cells with intact GVs (bright inclusions, left) and following GV collapse by sonication (right). Scale bars: 20 µm. **(C)** Schematic overview of the RNA sequencing workflow. Cells with intact GVs were either left untreated or subjected to sonication to induce GV collapse, followed by RNA extraction and sequencing. **(D)** Volcano plot showing differential gene expression between collapsed and intact GV conditions. The x-axis represents log_2_ fold change (collapsed / control), and the y-axis shows statistical significance as –log_10_ (Benjamini-Hochberg adjusted p-value of the Wald test for the identified hits). Each dot corresponds to a transcript; significantly enriched or depleted genes (adjusted *p* < 0.05) are colored red or blue, respectively. Selected upregulated genes are labeled. Scientific illustrations are adapted from the NIAID NIH BioArt collection.

### Collapse of GVs in *E. coli* does not induce the expression of stress-responsive reporter genes

Recombinant expression of GVs in heterologous hosts such as *E. coli* is often a key step in developing biomedical applications, including the original development of acoustic reporter genes^6^. Motivated by the distinct transcriptional response to GV collapse observed in *D. flos-aquae*, we next asked whether a similar response could be detected for the collapse of these recombinantly expressed GVs in *E. coli*. Unlike *D. flos-aquae*, which lacks established genetic tools for plasmid transformation, *E. coli* offers the advantage of convenient plasmid-based gene expression. Therefore, rather than performing transcriptomic analysis, we opted to use reporter gene assays to examine the transcriptional response to GV collapse.

Specifically, we constructed plasmids that carried NanoLuc, a small luciferase enzyme for bioluminescence signals^38^, driven by stress-responsive promoters. We reasoned that bioluminescence would offer a high signal-to-background ratio and a rapid readout of GV-collapse-induced transcriptional activity. Initially, we designed plasmids placing NanoLuc under the control of *E. coli* promoters for *dnaK* and *clpXP*, which are known to be upregulated in response to protein misfolding^39^. These plasmids were co-transformed with the pST39-pNL29 plasmid encoding *B. megaterium*-derived pNL29 GVs (**Figure 2A**)^40^. After co-transforming the two plasmids into *E. coli*, GV expression was induced for at least 18 hours. Next, we collapsed the GVs using water bath sonication and measured NanoLuc luminescence as a proxy for promoter activation (**Figure 2B**). However, we observed no significant increase in luminescence in response to GV collapse, either compared to cells with intact GVs or to cells carrying a mRuby2-expressing control plasmid (“mRuby2 dummy”) (**Figure 2C**). As additional positive controls, we confirmed that the reporter constructs were functional by applying a non-lethal heat shock at the time of induction, which led to increased luminescence one hour later (**Figure 2D**). However, after 24 hours of culture—mirroring the GV expression timeline—the inducibility of the *pdnaK* reporter was diminished, whether paired with the GV operon or the control plasmid (**Figure 2E**). This suggested that cellular aging or high baseline promoter activity may blunt the response to additional stress stimuli.

**Fig 2.**
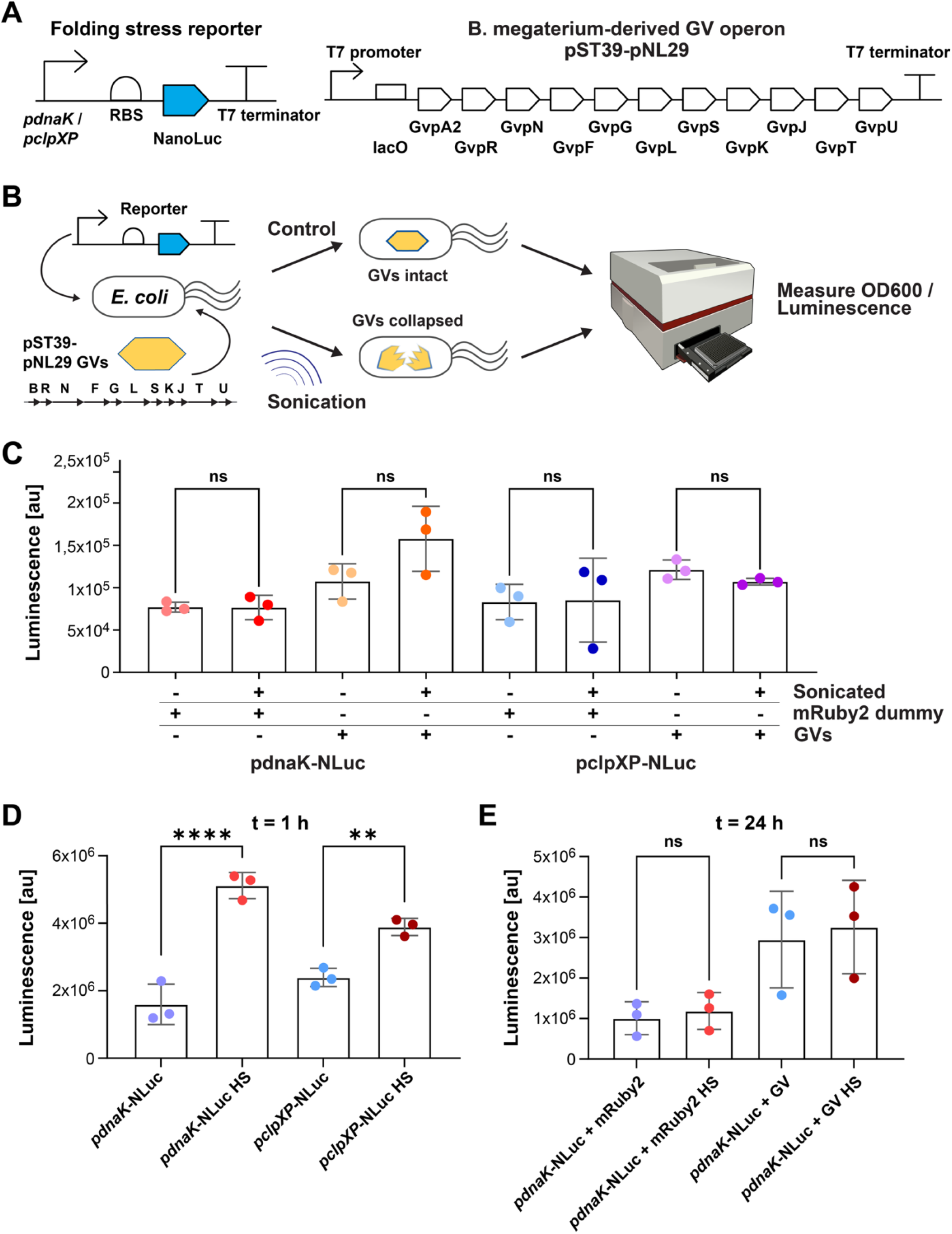
GV collapse does not induce a detectable protein folding stress response in *E. coli*. **(A)** SBOL diagrams of the constructs used: the left shows the NanoLuc-based protein folding stress reporters driven by heat shock promoters (*pdnaK* or *pclpXP*); the right shows the pNL29 GV expression operon (*pST39-pNL29*) under T7 promoter control, inducible with IPTG. **(B)** Schematic of the GV collapse assay in *E. coli*. Cells co-transformed with the reporter and GV plasmids were induced with IPTG to express GVs, then either sonicated to collapse GVs or left untreated. NanoLuc luminescence was measured to assess stress response. **(C)** Bar graph showing NanoLuc expression following GV collapse. Y-axis: luminescence normalized by OD_600_; X-axis: combinations of GV or control plasmid that express mRuby2 (mRuby2 dummy) with either reporter. Triplicate samples; gain = 212. No significant differences were observed (ANOVA with Brown-Forsythe correction; p = 1, 0.7, 1, 1). **(D)** Heat shock serves as a positive control for the reporter functionality. Reporter constructs were tested under 42 °C for 1 hour. Significant induction was observed between heat shocked (indicated by “HS”) and control samples (*pdnaK*: p = 0.0001; *pclpXP*: p = 0.0036; gain = 144). **(E)** Testing GV-expressing cells for heat shock response confirms reporter function is not impaired by GV co-expression. No significant differences (p = 0.96, 0.9; gain = 150). Illustrations adapted from the NIAID NIH BioArt collection.

To improve the sensitivity and dynamic range of the reporters, we designed two additional constructs following published strategies for enhanced stress sensing^41^. First, we combined the *pdnaK* reporter with the transcriptional regulator HspR and HspR-associated inverted repeats^42^, so that expression of the reporter is more stringently controlled by cellular stress (**Figure 3A**). Second, we implemented a dual-layer, two *pibpA* promoters riboswitch system using a cis-repressing RNA upstream of the NanoLuc ribosome binding site, which would be relieved by a trans-activating RNA expressed from the second *pibpA* unit^43^ (**Figure 3B**). Repeating the GV collapse experiments using either this riboswitch-based construct or the HspR-regulated construct, we again observed no significant luminescence change in response to GV collapse (**Figures 3C-D**). However, both constructs responded to heat shock in a similar fashion as the initial reporters (**Figures 3E–H**).

**Fig 3.**
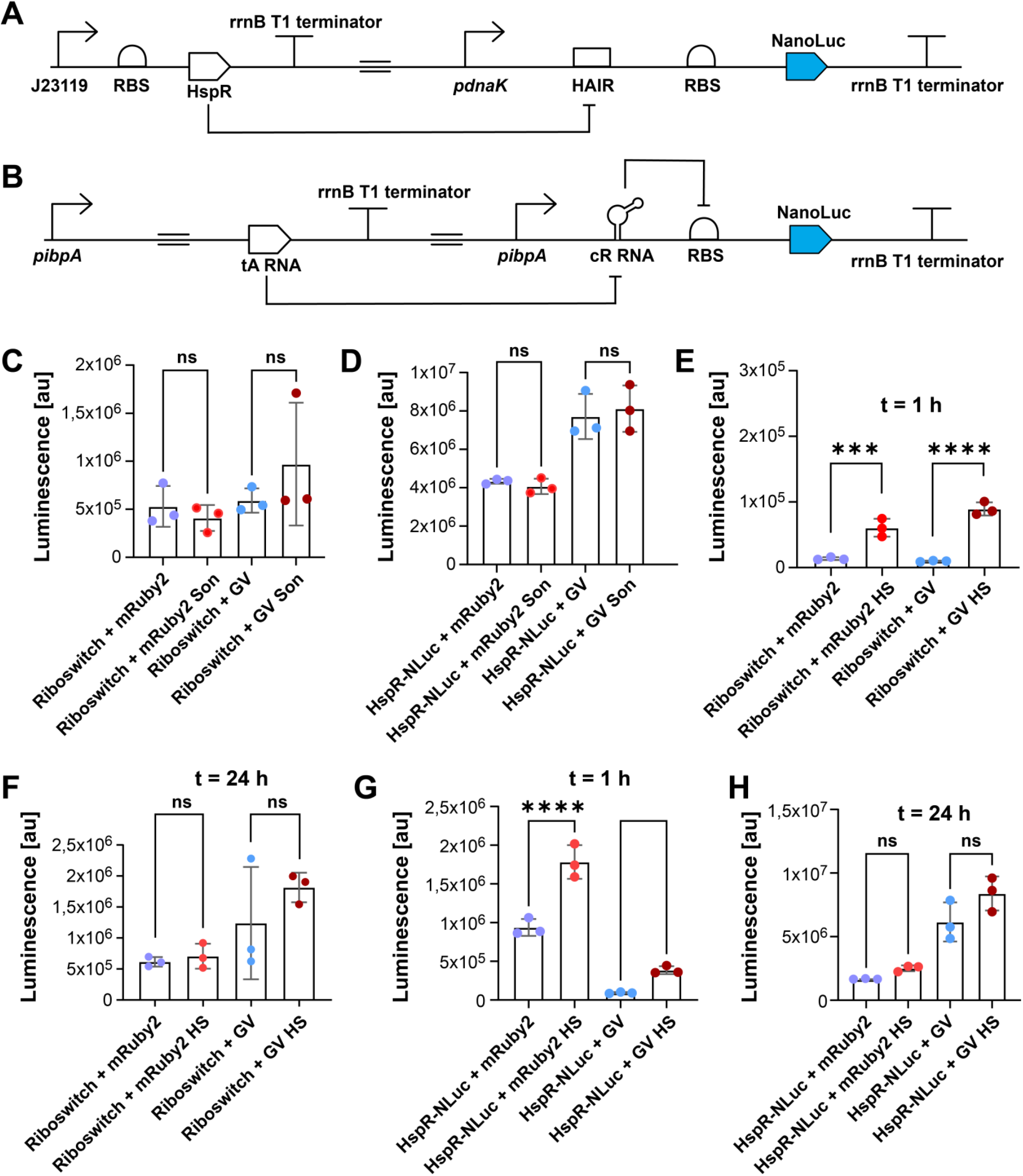
Optimized reporter circuits do not detect GV collapse in *E. coli*. **(A)** SBOL diagram of the HspR-based reporter circuit. HspR is constitutively expressed and represses the downstream *pdnaK*-NanoLuc expression via interaction with the HAIR element. Heat stress or unbound DnaK diminishes HspR repression, enabling reporter expression. **(B)** SBOL diagram of the riboswitch-based reporter circuit. A trans-activating (tA) RNA is under control of the pibpA promoter. Downstream, the pibpA promoter drives the expression of NanoLuc, which is inhibited by a cis-repressing (cR) RNA. The tA RNA binds the cR RNA to resolve the repressing effect. (**C**) Bar graph indicating the results of the GV-collapse induced protein folding stress assay. The gain was set to 150. The Y-axis indicates the luminescence measured divided by the OD_600_ measured for each assayed sample in triplicates. The plasmids used per condition are indicated on the x-axis: the Riboswitch construct with either the mRuby2 dummy plasmid or the pST39-pNL29 gas vesicle plasmid. Samples are marked as either sonicated or a control condition. Results of one-way Anova followed by Brown-Forsythe testing are indicated. P-values (left to right) are 1, 0.98. (**D**) Bar graph indicating the results of the GV-collapse induced protein folding stress assay. The gain was set to 150. The Y-axis indicates the luminescence measured divided by the OD_600_ measured for each assayed sample in triplicates. The plasmids used per condition are indicated on the x-axis: the HspR construct with either the mRuby2 dummy plasmid or the pST39-pNL29 gas vesicle plasmid. Samples are marked as either sonicated or a control condition. Results of one-way Anova followed by Brown-Forsythe testing are indicated. P-values (left to right) are 1, 0.97. (**E**) Bar graph indicating the results of the heat-shock induced protein folding stress assay measured one hour after the addition of IPTG. The Y-axis indicates the luminescence measured divided by the OD_600_ measured for each assayed sample in triplicates. The gain was set to 150. The X-axis indicates the plasmids used for transformation: GVs used are derived from the pST39-pNL29 plasmid, the mRuby2 dummy expresses mRuby2 under control of the T7 promoter and IPTG instead, and the reporter used was the Riboswitch construct. Samples are marked as either heat-shocked (“HS”) or a control condition. Results of one-way Anova followed by Brown-Forsythe testing are indicated. P-values (left to right) are 0.0003, and 0.0001. (**F**) As (E), but the experiment was performed 24 hours after adding IPTG. Results of one-way Anova followed by Brown-Forsythe testing are indicated. P-values (left to right) are 1, 0.77. (**G**) Bar graph indicating the results of the heat-shock induced protein folding stress assay measured one hour after the addition of IPTG. The Y-axis indicates the luminescence measured divided by the OD_600_ measured for each assayed sample in triplicates. The gain was set to 150. The X-axis indicates the plasmids used for transformation: GVs used are derived from the pST39-pNL29 plasmid, the mRuby2 dummy expresses mRuby2 under control of the T7 promoter and IPTG instead, and the reporter used was the HspR construct. Samples are marked as either heat-shocked (“HS”) or a control condition. Results of one-way Anova followed by Brown-Forsythe testing are indicated. P-values (left to right) are 0.0001, and 0.04. (**H**) As (G), but the experiment was performed 24 hours after adding IPTG. Results of one-way Anova followed by Brown-Forsythe testing are indicated. P-values (left to right) are 0.54, 0.06.

One possible explanation for the absence of molecular chaperone upregulation is that the transcriptional response in *E. coli*, if it exists, is too subtle to be detected by our current reporter systems. For example, the corresponding transcripts in *D. flos-aquae* were upregulated between 3- and 10-fold, changes that may be below the detection threshold of our assay. Additionally, differences in expression dynamics between the two species may play a role: the time frame and consistency of GV expression in *E. coli* may not mirror that of the native host. It is also conceivable that the magnitude of protein folding stress induced by GV collapse in *D. flos-aquae* depends on uniformly high levels of GV expression across the population, a condition that may not be met in heterologous systems such as *E. coli*^44^. Finally, it remains possible that *D. flos-aquae* possesses a specialized mechanism for sensing GV collapse, one that is absent in heterologous hosts.

### Search for specific *E. coli* proteins that bind collapsed gas vesicles

Next, we asked whether specific *E. coli* proteins exhibit preferential affinity for collapsed, but not intact, GVs. To test this, we developed a pull-down assay using collapsed GVs as bait to capture *E. coli* proteins (**Figure 4A**). GVs purified from *D. flos-aquae* were surface-biotinylated and immobilized on streptavidin-coated magnetic beads. Successful immobilization was confirmed by the flotation of these magnetic beads ( **Figure 4B**). We then either sonicated the beads to collapse the GVs or left them untreated as a control before incubating them with *E. coli* lysates. After incubation and washing, any differential protein binding between collapsed and intact GVs would suggest selective interactions. Samples were analyzed by LC-MS/MS in triplicate to identify proteins associated with the beads.

**Fig 4.**
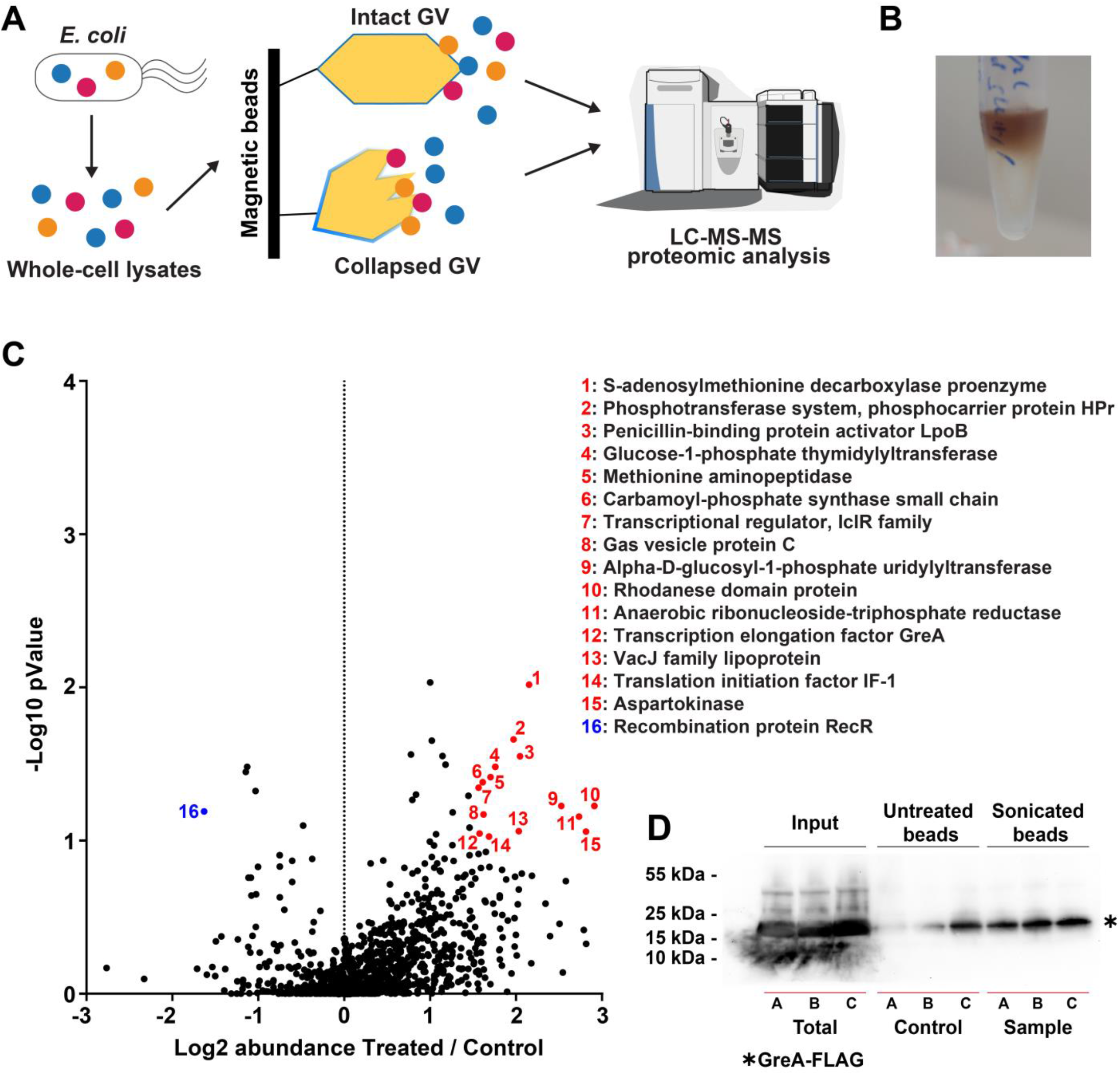
*E. coli* proteins do not exhibit selective binding to collapsed GVs. **(A)** Schematic of the pull-down assay. *E. coli* lysates were incubated with streptavidin-coated magnetic beads loaded with biotinylated GVs to create GV-affinity beads. Beads were either left intact or sonicated to collapse the GVs. Following washing, LC-MS/MS was used to identify proteins that differentially bind to collapsed versus intact GVs. **(B)** Photograph showing buoyant GV-coated magnetic beads after incubation with biotinylated GVs, demonstrating successful loading. **(C)** Volcano plot of LC-MS/MS results comparing proteins bound to collapsed versus intact GVs. The x-axis shows log_2_ fold change (collapsed / intact) based on label-free quantification, averaged across biological triplicates. The y-axis shows statistical confidence as –log_10_ (p-value). Spheres indicate the individual proteins identified, and those proteins identified as enriched or depleted in the collapsed GV fraction are marked in red and blue, respectively, with protein names given. **(D)** Western blot image detecting GreA-FLAG in pull-down fractions. *E. coli* whole-cell lysates (“Input”), and material eluted from intact (“Untreated beads”) or collapsed (“Sonicated beads”) GV-coated beads were probed with anti-FLAG antibody. Molecular weight markers are shown on the left. A–C correspond to replicate samples. Quantification of band intensity is summarized in Supplementary Table 5. Image processing and quantification were performed in Fiji^54^. Illustrations adapted from the NIAID NIH BioArt collection.

A broad range of *E. coli* proteins was recovered (**Supplementary Table 2** and **Figure 4C**); however, based on their annotated functions, we consider most of them unlikely to serve as binders for large protein assemblies such as collapsed GVs, with the possible exception of GreA, a transcriptional regulator (#12 in **Figure 4C**). Given the possibility that recruitment of a transcription factor could be related to a GV-triggered transcriptional response, we chose to further investigate GreA. *E. coli* cells were transformed with a plasmid expressing GreA fused to a C-terminal FLAG tag, and the pull-down assay was repeated using lysates from these cells. Western blotting using anti-FLAG antibodies revealed approximately twofold enrichment of GreA in the collapsed GV condition relative to controls (**Figure 4D**), normalized by total protein staining (**Supplementary Figure 1A**; quantification in **Supplementary Table 5**). This modest enrichment suggests that GreA association with collapsed GVs is weak and likely not indicative of a specific or functional interaction.

### Search for endogenous *D. flos-aquae* proteins that bind collapsed gas vesicles

To explore whether *D. flos-aquae* harbors specific proteins that interact preferentially with collapsed GVs and could contribute to a transcriptional response, we performed a similar pull-down assay as shown in Figure 4A. Magnetic beads coated with GVs were prepared as described above, then either left intact or collapsed by water bath sonication. Each condition was incubated in triplicate with cleared whole-cell lysates of *D. flos-aquae* instead of the lysates of *E. coli*. Following incubation and elution, the bound proteins were analyzed by LC-MS/MS using label-free quantification. We identified a number of proteins that were moderately enriched in the collapsed GV samples, but their fold changes in abundance compared to the control condition were modest (**Figure 5A** and **Supplementary Table 3**).

**Fig 5.**
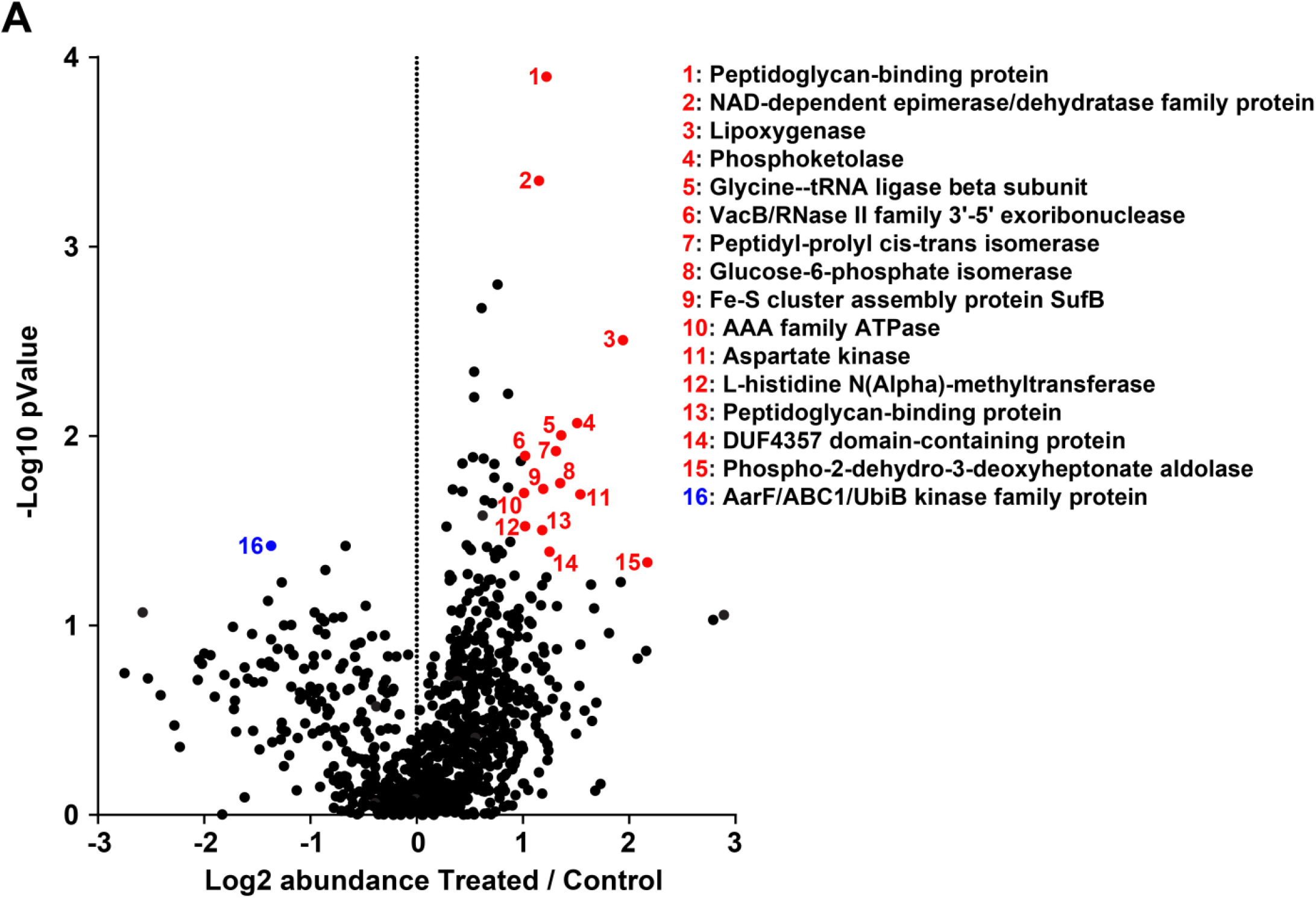
A limited number of *D. flos-aquae* proteins exhibit preferential binding to collapsed GVs. **(A)** Schematic of lysates from *D. flos-aquae* were incubated with streptavidin-coated magnetic beads loaded with biotinylated GVs. Beads were either sonicated to collapse the GVs or left intact as controls. After incubation and washing, bound proteins were analyzed by LC-MS/MS to identify candidates with differential affinity for collapsed GVs. **(B)** Volcano plot showing LC-MS/MS results comparing protein binding to collapsed versus intact GVs in *D. flos-aquae*. The x-axis represents log_2_ (fold change) in label-free quantification between collapsed and intact GV conditions, averaged over triplicates. The y-axis displays statistical significance as –log_10_ (p-value). Individual proteins are shown as points; those significantly enriched or depleted in the collapsed GV condition are highlighted in red or blue, respectively, with annotated protein names. Illustrations adapted from the NIAID NIH BioArt collection.

Compared to the *E. coli* dataset, *D. flos-aquae* exhibited a greater number of statistically significant differential binders. However, the degree of binding preference still did not exceed a 4-fold difference. Proteins enriched in the collapsed GV condition included several metabolic enzymes, an exoribonuclease, an AAA+ ATPase, and a protein of unknown function. Interestingly, an AarF-like kinase was significantly depleted in the collapsed GV condition (**Supplementary Table 3**). These findings suggest that *D. flos-aquae*, like *E. coli*, does not appear to possess a specific peptide or protein that directly reports on the structural status of GVs. It remains possible that the transcriptional response to GV collapse is mediated by proteins not detectable within the sensitivity range of this MS-based pull-down approach. Alternatively, the response may result from a biophysical change within the cell, such as altered turgor pressure or changes in free cytoplasmic volume, rather than from direct protein-level sensing of collapsed GVs.

## DISCUSSION

In this study, we tested the hypothesis that GV collapse may trigger a physiological response in host cells. Using transcriptomic and proteomic methods, we found that *D. flos-aquae* exhibits a distinct transcriptional response involving heat shock proteins, while *E. coli* showed some activity but lacked a coordinated upregulation of similar stress-related genes. In the second part of the study, we aimed to identify a molecular mechanism by which cells recognize collapsed GVs, potentially through proteins with selective binding. However, despite comprehensive proteomic analyses in both species, we did not identify strong candidate proteins that preferentially bind collapsed GVs.

In natural environments, studies suggest that *D. flos-aquae* cells may undergo GV collapse as part of their life cycle. Metabolite accumulation during photosynthesis or the influx of potassium ions in conjunction with light exposure can increase intracellular turgor pressure, potentially leading to GV collapse ^45, 46^. This process allows cells to descend to deeper water layers, regenerate GVs, and then return to the surface to resume photosynthesis, enabling dynamic vertical migration for optimal light exposure^47^. It is therefore plausible that native GV-producing organisms have evolved mechanisms to detect and respond to GV collapse.

From a biophysical perspective, collapsed GVs may resemble protein aggregates, which cells often recognize and respond to. Supporting this idea, our RNAseq data from *D. flos-aquae* show significant upregulation of molecular chaperones following GV collapse. Notably, *clpB, dnaK*, and a J-protein family member (*HGD76_RS05240*) were among the significantly induced transcripts, consistent with the known function of the bacterial protein disaggregation machinery^48^ powered by Hsp100 family proteins^49, 50^. Interestingly, the *D. flos-aquae* genome encodes two homologs of ClpB, ClpBI and ClpBII, a feature also observed in other cyanobacteria^37^. While not all cyanobacteria express GVs, the presence of multiple ClpB homologs may support cellular mechanisms to handle GV collapse in native hosts. In addition to molecular chaperones, we observed upregulation of other genes, including a DMT family transporter, an ion channel protein (*fluC*), an enzyme involved in molybdopterin biosynthesis (*moaE*), and several hypothetical proteins. The functions of these genes are less clear but may relate to the broader metabolic or membrane-associated consequences of GV collapse rather than direct involvement in disaggregation.

Although we did not identify a definitive molecular sensor or binding partner for collapsed GVs, our study provides a foundational view of GV-related cellular responses through an omics-based lens. The integration of transcriptomic and proteomic data enabled a broad exploration of both gene expression changes and protein interactions in response to GV collapse in native and heterologous contexts. While our results did not yield strong candidate effectors, they revealed several protein hits that warrant further focused functional investigation and underscore the need for more sensitive tools to probe GV-related signaling. We hope this study contributes to advancing the understanding of GV biophysics and cellular microbiology, and motivates future mechanistic insights into how cells detect and manage large intracellular protein aggregates, including collapsed gas vesicles.

## METHODS

### Plasmid construction

To generate *pdnaK*-NanoLuc and *pclpXP* plasmids, PCR was performed to amplify the *pdnaK-NanoLuc* and *pclpXP-NanoLuc* fragments from synthetic genes and insert them into a plasmid backbone with spectinomycin resistance markers and a CloDF13 origin of replication^51^ via Gibson assembly to be co-expressible with the pST39-pNL29 GV plasmid using Q5 Hot Start High-Fidelity DNA polymerase and HiFi DNA Assembly Master Mix (New England Biolabs (NEB), Ipswich, MA). To generate the *pdnaK*-NanoLuc plasmid with HspR and the HAIR element, first, the rrnB T1 terminator was amplified from a synthetic gene and inserted downstream of the *NanoLuc* gene in the *pdnaK-NanoLuc* plasmid. Subsequently, site-directed mutagenesis in combination with blunt end ligation was performed to insert the *HAIR* element upstream of the ribosomal binding site (RBS) using KLD Enzyme Mix (NEB, Ipswich, MA). Finally, an expression unit *J23119-RBS-HspR-rrnB T1* was amplified from a synthetic gene and inserted into this plasmid *via* Gibson assembly. To create the Riboswitch plasmid construct, a synthetic gene element *pibpA-cis-repressing RNA-loop-RBS* was inserted upstream of the NanoLuc gene in the *pdnaK-NanoLuc-rrnB T1* plasmid *via* Gibson assembly. Subsequently, a second synthetic expression unit *pibpA-TAAT-trans-activating-RNA-rrnB T1* was inserted upstream of this element in the same fashion. To express GreA-FLAG, a synthetic element *greA* was amplified and inserted into a plasmid backbone with the p15A origin of replication and a Kanamycin resistance marker for expression ^51^, followed by mutagenesis as above to install a C-terminal FLAG-tag.

Supplementary Table 4 contains a list of DNA elements and oligonucleotides used. Supplementary Figure 2 contains visual representations of the plasmids generated. Primers were designed and representative plasmid maps generated using SnapGene (GSL Biotech LLC, San Diego, USA).

### Transformation of *E. coli*

For cloning purposes, plasmids created by Gibson assembly or site-directed mutagenesis were incubated with competent NEB Turbo *E. coli* cells for 30 minutes on ice before transformation by heat shock for 30 seconds at 42 °C in a water bath. Cells were left to recover on ice for 5 minutes before the addition of 200 µL SOC media in a 1.5 mL tube and transformants were isolated on selective agar plates with either 50 µg/mL Kanamycin or 75 µg/mL Spectinomycin. Plasmid sequences were confirmed by Sanger sequencing (Eurofins Scientific, Luxembourg, Luxembourg). For experiments expressing GVs, reporter genes or GreA-FLAG, competent *E. coli* BL21 (DE3) cells were transformed with the indicated plasmids and selected for with the same antibiotics or 100 µg/µL Carbenicillin for the pST39-pNL29 GV plasmid. For GV plasmids, 1 % (w/v) glucose was included during growth to minimize background expression of GV proteins.

### Culturing of *D. flos-aquae* and RNA extraction

*D. flos-aquae* cultures were grown in BG11-media (Sigma-Aldrich, St. Louis, MO, USA) under 1% CO_2_ at 26 °C and approximately 50 µM photons / m^2^ / s^-1^ from a white LED light source. Three separate cultures of 50 mL cells each were collected at an OD_600_ of 2.0 and split into two samples of 25 mL each. Control samples were left at room temperature, GV collapse samples were subjected to waterbath sonication in a Bransonic CPHX Digital Bath 8800 (Emerson, St. Louis, MO, USA) for three sets of 10 seconds each. All samples were then collected on Whatman filter membranes (Cytiva, Marlborough, USA) and resuspended in 5 mL phosphate-buffered saline before addition of 10 mL RNAProtect reagent (Qiagen, Hilden, Germany). After agitating the samples for 5 seconds, they were incubated for 5 minutes, collected by vacuum filtration onto Whatman filter membranes (Cytiva, Marlborough, USA) and flash frozen in liquid nitrogen for subsequent use.

RNA extraction was performed with a Qiagen RNeasy Mini kit (Qiagen, Hilden, Germany) following the manufacturer’s instructions, with modifications to the lysis procedure as follows: Lysozyme was added to 2.5 mg / mL and supplemented with 40 µM Spermine as well as 0.2% of the detergent C7Bz0 and 2% of the detergent SB3-14, in addition to 50 mM dithiothreitol^52^. Cells were then subjected to homogenization in a Bead Mill Homogenizer (VWR, Radnor,PA, USA) for 2 cycles with 2.4 mm metal beads (VWR, Radnor,PA, USA) and 3 cycles with 0.5 mm glass beads (VWR, Radnor,PA, USA) for 45 seconds each while resting on ice in between steps.

### RNA sequencing

Samples were submitted to Azenta Life Sciences (Burlington, USA) for RNA sequencing. RNA samples were quantified using a Qubit 2.0 Fluorometer (ThermoFisher Scientific, Waltham, MA, USA) and RNA integrity was checked with a 4200 TapeStation (Agilent Technologies, Palo Alto, CA, USA).

rRNA depletion sequencing library was prepared by using three probes from QIAGEN FastSelect rRNA 5S/16S/23S Kit (Qiagen, Hilden, Germany), respectively. RNA sequencing library preparation was performed using the NEBNext Ultra II RNA Library Preparation Kit for Illumina, following the manufacturer’s recommendations (NEB, Ipswich, MA, USA). In brief, enriched RNAs were fragmented for 15 minutes at 94 °C. First strand and second strand cDNA were subsequently synthesized. cDNA fragments were end repaired and adenylated at the 3’ends, and universal adapters were ligated to cDNA fragments, followed by index addition and library enrichment with limited cycle PCR. Sequencing libraries were validated using the Agilent Tapestation 4200 (Agilent Technologies, Palo Alto, CA, USA), and quantified using the Qubit 2.0 Fluorometer (ThermoFisher Scientific, Waltham, MA, USA) as well as by quantitative PCR (KAPA Biosystems, Wilmington, MA, USA).

The sequencing libraries were multiplexed and clustered onto 1 lane of a flowcell. After clustering, the flowcell was loaded onto the Illumina HiSeq 4000 instrument according to the manufacturer’s instructions. The samples were sequenced using a 2×150bp Paired End (PE) configuration. Image analysis and base calling were conducted by the Illumina Control Software. Raw sequence data (.bcl files) generated were converted into fastq files and de-multiplexed using Illumina bcl2fastq 2.17 software. One mismatch was allowed for index sequence identification.

### Assay of *in vivo* stress response for *E. coli*

Cells co-transformed with the reporter plasmids and either the pST39-pNL29 plasmid or the mRuby2 dummy plasmid were plated onto LB-agar plates with 75 µg/mL Spectinomycin, 100 µg/µL Carbenicillin and 1 % (w/v) glucose. Individual colonies from the different assay conditions were then grown in 1 mL LB media with the same antibiotics and glucose overnight. The next day, all cells were diluted 50-fold into 3 mL of LB media with the same antibiotics and 0.2 % (w/v) glucose and grown at 37 °C while shaking at 600 rpm until the OD_600_ exceeded 0.3 for all cultures. At this point, IPTG was added for induction of GV expression to a concentration of 20 µM and the temperature of incubation was reduced to 30 °C.

To assay heat shock response, 200 µL of cultures were transferred to 1.5 mL reaction vessels and subjected to either 30 °C or 42 °C for one hour either at the timepoint of induction, or at the timepoint cells were subjected to sonication. Cells sampled at the later timepoint were diluted 10-fold in LB media before measurement. To assay the GV collapse response, 1.6 mL of cells were sampled 24 hours after addition of IPTG, transferred to 2 mL reaction vessels and centrifuged at 400 x g for 1 hour. Then, to ensure that cells containing GVs were sampled, 200 µL of the top natant of these samples were transferred to 1.5 mL reaction vessels and subjected to waterbath sonication in a Bransonic CPHX Digital Bath 8800 (Emerson, St. Louis, MO, USA) for three sets of 10 seconds and returned to incubate at 30 °C for 1 hour.

To quantify the expression of NanoLuc, luminescence was measured. First, 200 µL of cells were transferred to a clear 96-well plate (#3370, Corning, Corning, NY, USA) to take OD_600_ measurements in a Biotek Synergy H4 Multimode (Agilent, Santa Clara, CA, USA) plate reader. Cells were then subsampled and 10 µL of each sample were mixed with 10 µL of prepared NanoGlo Live Cell Substrate (Promega, Madison, WI, USA) and 30 µL of LB media in black opaque 96-well half-area plates (#3694, Corning, Corning, NY, USA) before measuring luminescence in 8 repeat measurements at a read height of 4.5 mm without an emission filter.

The results of measurements were reported as arbitrary units [au] and for each experiment, the gain used on the plate reader is indicated. To normalize the luminescence signals to the quantity of cells assayed per condition, the measured luminescence is divided by the OD_600_ value measured for each sample.

### Preparation of GV-loaded magnetic beads

To prepare magnetic beads loaded with GVs, GVs were first prepared as described previously ^40^ and adjusted to a volume of 500 µL at an OD_600_ of 4 in a 1.5 mL reaction vessel. *EZ-Link*-Sulfo-N-hydroxy-succinimide-LC-Biotin (Thermo Fisher Scientific, Waltham, MA, USA) was then added to a concentration of 20 µM and incubated while agitating for at least 4 hours at 4 °C to covalently link biotin to the surface of the GVs. GVs were then dialyzed against 1 liter of PBS for 4 hours followed by a second dialysis step over night at 4 °C.

Pierce Streptavidin-linked magnetic beads (Thermo Fisher Scientific, Waltham, MA, USA) were washed with bind/wash buffer (30 mM HEPES, pH 7.4, 150 mM NaCl and 0.1% Tween 20) once before adding the biotinylated GVs to incubate at 4 °C overnight while rolling. The GV-coated beads were then washed with bind/wash buffer followed by PBS before being used for the next experiment. Successful loading with GVs was confirmed by observing buoyancy of the beads after one day in storage.

### Pull-down assay with bacterial lysates

Pull-downs were performed in triplicates, using 50 µL of GV-loaded magnetic beads per condition. First, cell lysates were prepared, using lysis buffer (30 mM HEPES, pH 7.4, 150 mM NaCl, 5 mM MgCl_2_) and a Branson Sonifier 250 probe sonicator (Emerson, St. Louis, MO, USA) for *E. coli* or the lysis method described above for *D. flos-aquae*. Each replicate was prepared from a total of 50 mL cells which had been collected by centrifugation for *E. coli*, or filter membrane for *D. flos-aquae* and buffers were supplemented with 1 mM dithiothreitol, 2.5 mg / mL Lysozyme, 100 µg / mL DNaseI (Sigma-Aldrich, St. Louis, MO, USA), as well as a protease inhibitor cocktail with Leupeptin (500 ng / mL), Pepstatin (700 ng / mL), E-64 (3.5 µg / mL) and PMSF (4 µg / mL). Lysates were cleared by centrifugation at 3000 x g for 5 minutes and supernatants were recovered. For each sample, the beads were split into two groups where one was subjected to waterbath sonication in 1.5 mL reaction vessels for three cycles of 10 seconds each. Beads were then mixed with 500 µL of the lysates and incubated while rolling at 4 °C overnight, for a total of three control samples and three samples with collapsed GVs. After incubation, beads were washed with TBST three times and then resuspended in 50 µL of H_2_O before proceeding with LC-MS-MS analysis.

### LC-MS-MS analysis

#### Sample digestion

The samples were prepared similar to as described before^53^. Briefly, 25 µg of sample were solubilized with 40 µL of 5% SDS, 50mM TEAB, pH 7.55 and incubated at RT for 30 minutes. The supernatant containing proteins of interest was then transferred to a new tube and reduced by making the solution 10mM TCEP (Thermo, #77720) and incubated at 65°C for 10min. The sample is then cooled to RT and 3.75 uL of 1M iodoacetamide acid added and allowed to react for 20 minutes in the dark after which 0.5uL of 2M DTT is added to quench the reaction. 5 uL of 12% phosphoric acid is added to the 50uL protein solution. 350uL of binding buffer (90% Methanol, 100mM TEAB final; pH 7.1) is then added to the solution. The resulting solution is added to S-Trap spin column (protifi.com) and passed through the column using a bench top centrifuge (30s spin at 4,000g). The spin column is washed with 400uL of binding buffer and centrifuged. This is repeated three times. Trypsin is added to the protein mixture in a ratio of 1:25 in 50 mM TEAB, pH = 8, and incubated at 37^○^C for 4 hours. Peptides were eluted with 80uL of 50mM TEAB, followed by 80uL of 0.2% formic acid, and finally 80 uL of 50% acetonitrile, 0.2% formic acid. The combined peptide solution is then dried in a speed vac and resuspended in 2% acetonitrile, 0.1% formic acid, 97.9% water and placed in an autosampler vial.

#### NanoLC MS/MS Analysis

Peptide mixtures were analyzed by nanoflow liquid chromatography-tandem mass spectrometry (nanoLC-MS/MS) using a nano-LC chromatography system (UltiMate 3000 RSLCnano, Dionex), coupled on-line to a Thermo Orbitrap Fusion mass spectrometer through a nanospray ion source. A trap and elute method was used. The trap column was a C18 PepMap100 (300um X 5mm, 5um particle size) from ThermoScientific The analytical columns was an Acclaim PepMap 100 (75um X 25 cm) from (Thermo Scientific). After equilibrating the column in 98% solvent A (0.1% formic acid in water) and 2% solvent B (0.1% formic acid in acetonitrile (ACN)), the samples (2 µL in solvent A) were injected onto the trap column and subsequently eluted (400 nL/min) by gradient elution onto the C18 column as follows: isocratic at 2% B, 0-5 min; 2% to 24% B, 5-86 min; 24% to 44% B, 86-93 min; 44% to 90% B, 93-95 min; 90% B for 1 minute, 90% to 10% B, 96-98 min; 10% B for 1 minute 10% to 90% B, 99-102 min 90% to 4% B; 90% B for 3 minutes; 90% to 2%, 105-107 min; and isocratic at 2% B, till 120 min.

LC-MS/MS data were acquired using XCalibur, version 2.5 (Thermo Fisher Scientific) in positive ion mode using a top speed data-dependent acquisition (DDA) method with a 3 sec cycle time. The survey scans (*m/z* 350-1500) were acquired in the Orbitrap at 120,000 resolution (at *m/z* = 400) in profile mode, with a maximum injection time of 100 msec and an AGC target of 400,000 ions. The S-lens RF level is set to 60. Isolation is performed in the quadrupole with a 1.6 Da isolation window, and CID MS/MS acquisition is performed in profile mode using rapid scan rate with detection in the ion-trap, with the following settings: parent threshold = 5,000; collision energy = 32%; maximum injection time 56 msec; AGC target 500,000 ions. Monoisotopic precursor selection (MIPS) and charge state filtering were on, with charge states 2-6 included. Dynamic exclusion is used to remove selected precursor ions, with a +/-10 ppm mass tolerance, for 15 sec after acquisition of one MS/MS spectrum.

#### Database Searching

Tandem mass spectra were extracted and charge state deconvoluted by Proteome Discoverer (Thermo Fisher, version 2.2.0388). Charge state deconvolution and deisotoping were not performed. All MS/MS samples were analyzed using Sequest (Thermo Fisher Scientific, San Jose, CA, USA; version Minora in Proteome Discoverer 2.5.0.402). Sequest was set up to search *Dolichospermum flos-aquae* and a contaminant cRAP_formatted.fasta with the digestion enzyme trypsin. Sequest was searched with a fragment ion mass tolerance of 0.60 Da and a parent ion tolerance of 10.0 PPM. Carbamidomethyl of cysteine was specified in Sequest as a fixed modification. Deamidated of asparagine and glutamine, oxidation of methionine and acetyl of the n-terminus were specified as variable modifications.

### Phase-contrast imaging

Phase contrast imaging was performed by transferring 5 µL of bacterial cultures to glass object carriers and adding a drop of Fluoroshield Mounting Medium (Sigma-Aldrich, St. Louis, MO, USA). Drops were mixed with a pipette tip before a coverslip was added and cells were left to interact with the mounting media for at least 24 hours. An Eclipse TI2 inverted microscope (Nikon, Melville, NY, USA) was used to image cells, with a 40x phase contrast objective a 1.5x tube lens and a Ph2 condenser annulus. Exposure time was kept at 70 milliseconds. Phase contrast images were analyzed and presented using *Fiji*^54^.

### Western blotting and immunostaining

To prepare Western blotting, of GreA-FLAG samples isolated with GV-coated magnetic beads, 30 µL of the beads were mixed with 30 µL of 2x Laemmli buffer and incubated at 98 °C for 10 minutes in 1.5 mL reaction vessels. Total protein lysates of E. coli were sampled analogously. Samples were then spun at 17 000 xg for 1 minute to remove the beads and 25 µL of each sample were subjected to SDS-PAGE using 4-20% Mini-PROTEAN TGX precast protein gels (Bio-Rad, Hercules, CA, USA). 5 µL PageRuler Plus (Thermo Fisher Scientific, Waltham, MA, USA) were used as a marker of molecular weight.

After SDS-PAGE, Western blotting was performed with a Trans-Blot Turbo system and Trans-Blot Turbo Mini 0.2 µm Transfer-Packs (Bio-Rad, Hercules, CA, USA), choosing the “Standard” setting of the instrument. To assay total protein as a loading control, staining of the PVDF membrane was carried out using Revert 700 Total Protein Stain (LI-COR, Lincoln, NE; USA) following the manufacturer’s instructions before detection with a FluorChem M imager (Protein Simple, San Jose, CA, USA) using the “Multifluor Red” setting. The membrane was then transferred to a solution of 5% fat-free milk powder (w/v) in TBST (Tris-buffered saline with 0,1% Tween 20 detergent), before transferring it to a 1:500 dilution of rat anti-FLAG M2 antibodies (Agilent Technologies, Santa Clara, CA, USA) in TBST with 3 % milk powder (w/v) to be incubated overnight, rolling at 4 °C. A brief rinse in TBST was performed before transferring the membrane to a solution of goat anti-rat IgG (H+L) HRP-conjugated secondary antibodies (Invitrogen, Waltham, MA, USA) at a dilution of 1:5000 in 3 % milk + TBST. After incubating for 1 hour, the membrane was washed in TBST three times for 5 seconds, 5 minutes and 10 minutes respectively before a brief rinse in TBS and detection with Pierce ECL Western Blotting Substrate (Thermo Fisher Scientific, Waltham, MA, USA), choosing the “Chemi + Markers” setting and exposing for no more than 10 minutes. To prepare Western blot images for publication and quantify band sizes by pixel intensity, *Fiji* was used^54^.

### Statistical analysis

For Figure 2 and 3, one-way ANOVA testing was performed followed by Brown-Forsythe testing to compare the sonicated or heat-shocked versus the untreated condition. Statistical testing and data evaluation was carried out with *Excel* (*Microsoft*, Redmond, WA) and *Prism* (*Graphpad Software*, Boston, MA).

## ACKNOWLEDGMENTS

We thank the Shared Equipment Authority (SEA) at Rice University for the access to core facilities and instruments. This work was supported by the Cancer Prevention and Research Institute of Texas (CPRIT, RR190081), the NIH (R00EB024600, R21EB033607, and R35GM155015), the Welch Foundation (C-2249 and C-2069), G. Harold and Leila Y. Mathers Foundation (MF-2012-01314), John S. Dunn Foundation, and Rice University-Houston Methodist Seed Grant Program. M.I. acknowledges support from German Research Foundation (DFG) Postdoctoral Fellowship (511048568 IB 154/2-1). The UTMB Mass Spectrometry Facility is supported in part by Cancer Prevention Research Institute of Texas (CPRIT, RP190682).

## AUTHOR CONTRIBUTIONS

Conceptualization, G.J.L., M.I.; Methodology, G.J.L., M.I.; Investigation, M.I., A. H.; Formal Analysis, M.I.; Writing – Original Draft, G.J.L., M.I.; Supervision and Funding Acquisition, G.J.L..

## DATA AVAILABILITY

The set of plasmids generated for this study have been made available as Addgene deposit number 83964. The mass spectrometry proteomics data have been deposited to the ProteomeXchange Consortium via the PRIDE^55, 56^ partner repository with the dataset identifier PXD060779. The RNA sequencing data has been made available on NCBI GEO as GSE289028.

