## Supplemental Materials for "Transcriptomic and proteomic responses to gas vesicle collapse in native and engineered bacterial systems"

**Table of contents**

**Supplementary Table 1. Transcripts identified as differentially expressed by RNA sequencing.** The first sheet of this table lists all transcripts from *D. flos-aquae* that were identified as being differentially expressed after collapse of GVs by ultrasound. An explanation of the information given in each column is given in the second sheet.

**Supplementary Table 2. List of *E. coli* proteins identified by pull-down with GV-loaded magnetic beads**

The first sheet of this table lists all proteins identified by the pull-down experiment with GV-loaded magnetic beads and *E. coli* whole protein lysates. The rows listing likely false positives are greyed out. The second sheet lists the proteins that showed some degree of differential binding to beads with collapsed or intact GVs (with the –log(10) of the p-value greater than 1 and the log(2) of the fold-enrichment at least 1,5) and the third sheet provides an explanation of the information listed in the different columns.

**Supplementary Table 3. List of *D. flos-aquae* proteins identified by pull-down with GV-loaded magnetic beads**

The first sheet of this table lists all proteins identified by the pull-down experiment with GV-loaded magnetic beads and *D. flos-aquae* whole protein lysates. The second sheet lists the proteins that showed some degree of differential binding to beads with collapsed or intact GVs (with the –log(10) of the p-value greater than 1 and the log(2) of the fold-enrichment at least 1.5) and the third sheet provides an explanation of the information listed in the different columns.

**Supplementary Table 4. List of oligonucleotides and DNA elements used in this study.**

This table lists DNA elements used in this study as well as their source in the first sheet and oligonucleotides used in this study as well as their purpose in the second sheet.

**Supplementary Table 5. Quantified band intensity and calculations for GreA binding to collapsed GVs.**

This table lists the quantified band intensity of the GreA-FLAG bands from the western blot / immunostaining experiment shown in Supplementary Figure 3A and the total protein stain (as a loading control) shown in Supplementary Figure 3B. The relative abundance of GreA-FLAG in the sample with collapsed GVs is calculated and marked in red.

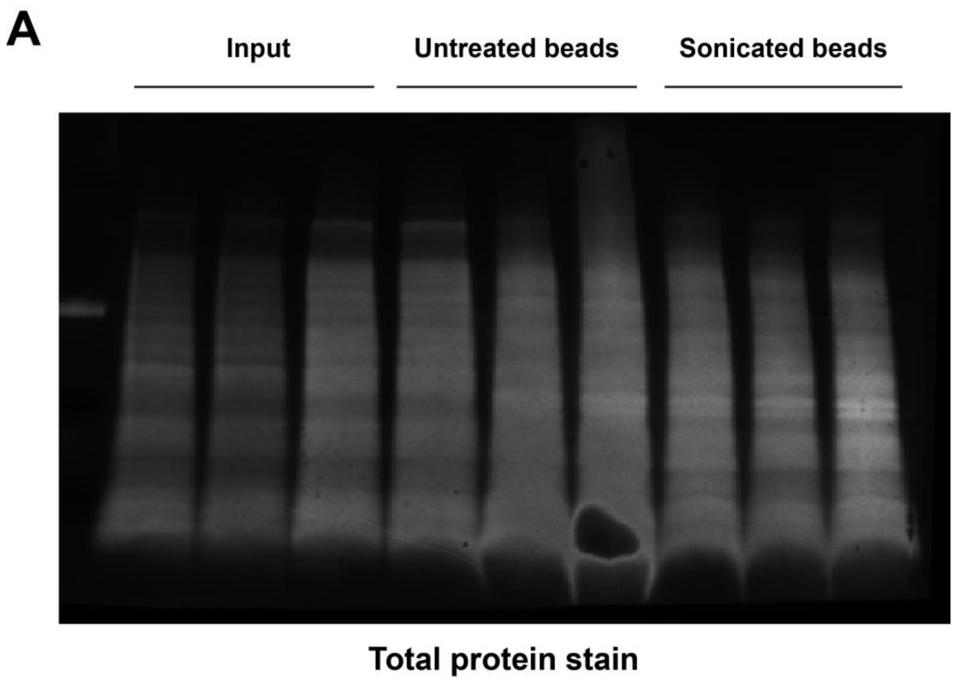


**Supplementary Figure 1. Immunostaining confirms a slight binding preference of GreA to collapsed GVs**

(**A**) An image of the western blot membrane shown in Figure 4D with a total protein stain applied as loading control. The setup of the samples is as in Figure 4D and the image was prepared and analyzed by *Fiji.*

A

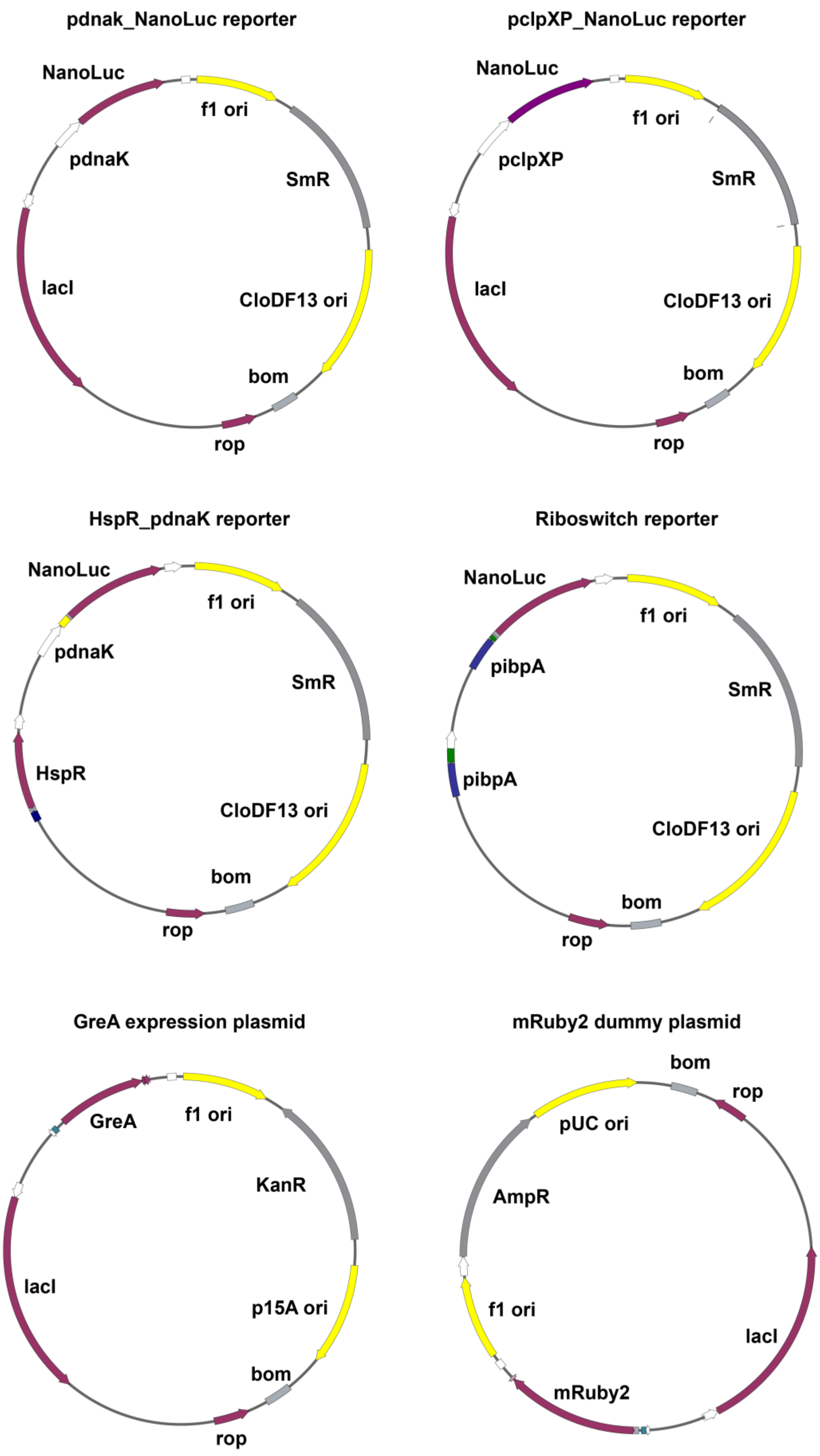
pdnak_NanoLuc reporter
NanoLuc
f1 ori
pdnaK
SmR
lacI
CloDF13 ori
bom
rop
pclpXP_NanoLuc reporter
NanoLuc
f1 ori
pclpXP
SmR
lacI
CloDF13 ori
bom
rop
HspR_pdnaK reporter
NanoLuc
f1 ori
pdnaK
SmR
HspR
CloDF13 ori
bom
rop
Riboswitch reporter
NanoLuc
f1 ori
pibpA
SmR
pibpA
CloDF13 ori
bom
rop
GreA expression plasmid
GreA
f1 ori
KanR
lacI
p15A ori
bom
rop
mRuby2 dummy plasmid
bom
rop
pUC ori
AmpR
f1 ori
lacI
mRuby2

**Supplementary Figure 2. Sketches of plasmid designs used for this study.**

This figure gives an overview of the plasmid architecture used for the *pdnaK*-NanoLuc reporter, the *pclpXP*-NanoLuc reporter, the mRuby2 dummy plasmid, the Riboswitch-NanoLuc reporter, the HspR-NanoLuc reporter as well as the plasmid used to drive expression of GreA-FLAG in *E. coli*. Elements indicated are the f1 origin of replication (f1 ori), the pUC origin of replication (pUC ori), the CloDF13 origin of replication (CloDF13 ori), the p15A origin of replication (p15A ori), the *rop* gene (rop), the *bom* gene (bom), a spectinomycin resistance marker (SmR), an ampicillin resistance marker (AmpR), the lacI expression cassette (lacI), the *NanoLuc* gene (NanoLuc), the *hspR* gene (HspR), the *greA-FLAG* gene (GreA), the *mRuby2* gene (mRuby2), and the promoters *pibpA*, *pdnaK*, *pclpXP.*
